# Model Choice Metrics to Optimize Profile-QSAR Performance

**DOI:** 10.1101/2022.08.22.504151

**Authors:** Stewart He, Sookyung Kim, Kevin S. McLoughlin, Hiranmayi Ranganathan, Da Shi, Jonathan E. Allen

## Abstract

**Background:** Predicting molecular activity against protein targets is difficult because of the paucity of experimental data. Approaches like multitask modeling and collaborative filtering seek to improve model accuracy by leveraging results from multiple targets, but are limited because different compounds are measured with different assays, leading to sparse data matrices. Profile-QSAR (pQSAR) 2.0 addresses this problem by fitting a series of partial least squares models for each target, using as features the predictions from single-task models on the remaining targets. This method has been shown to produce better results than single task and multitask models. However, the factors determining the success of pQSAR 2.0 have as yet not been characterized.

In this paper we examine the experimental conditions that lead to better pQSAR models. We limit the amount of data available to the method by retraining with decreasing amounts of data and explore the model’s ability to generalize to compounds that have never been assayed. Finally, we look at the properties of training data needed to demonstrate pQSAR improvement.

**Results:** We apply pQSAR 2.0 on a collection of GPCR and safety targets collected from Drug Target Commons, ExcapeDB, and ChEMBL. We found that pQSAR improved models on 34 of the 149 assays selected. In the other 115 assays, single task random forests offered better performance. There are many factors that contribute to an increase in performance, but the main factor is compound assay coverage. The pQSAR model improves when more compounds are measured in multiple assays.

**Conclusion:** It is necessary to consider the available data before applying pQSAR. Successful pQSAR models require a profile made of correlated targets that share compounds with other assays. This technique is best used when experimental data is available as random forest regressors often do not generalize well enough for virtual drug search applications.

## Background

Applying machine learning to property prediction in chemoinformatics presents many challenges. The key issue in chemoinformatics is not availability of compound structures or experimental data. Enamine’s REAL compound library [1] contains structures for over 13 billion readily synthesizable molecules. While large open-access experimental databases such as ChEMBL [2], Drug Target Commons (DTC) [3] and ExcapeDB [4] collect measurements from millions of physical and biochemical assays. The problem lies in the sparsity of the data and the need to develop predictive models that can generalize to new protein targets and new parts of chemical space.

As an example, the ChEMBL 28 release contains activity data for over 14,000 protein targets assayed against 2.1 million compounds. A dense matrix with targets along one axis and compounds on the other would take over 28 billion measurements to fill. At the time of publication only 17.3 million activities have been collected, many of which are replicate measurements; thus, less than 0.06% of the matrix is filled. The sparsity of data and the diversity of chemical structures tested vary considerably between protein targets. The vast size of chemical space makes it impossible to perform experimental measurements on a truly diverse sampling of chemical space. Thus, compound property prediction approaches must take into account this fundamental limitation in data coverage.

### Multitask Learning

Multitask learning is a machine learning approach that can be applied to learn protein-compound activity relationships across multiple targets and compound libraries simultaneously. In one example, a multitask model was trained against a fully filled matrix where every compound was tested against every target [5]. In this ideal case, the model demonstrated improvements in prediction accuracy over single task models. However, partially filled matrices are far more common given the expected scope of chemical space frequently interrogated. In another study, De et al. assembled a set of 243 PubChem assays using high-throughput screening fingerprints [6], obtaining a set of 454 kinase assays performed on 367 compounds [7]. The authors gradually deleted labels and measured the effect on performance. Smaller amounts of missing measurements were shown to not have a large impact on model performance. However, larger amounts of missing data could quickly degrade performance [8]. Multitask learning on the ChEMBL database is challenging due to the sparsity of most assays. Single task models frequently outperform multitask models with a large number of targets. Choosing targets by clustering on sequence similarity, using a method called multiple partial multi-task models, can boost performance of multitask models above single task models. [9].

### Collaborative Filtering

Collaborative filtering (CF) is one of the most successful approaches to predict missing values in sparse data matrices [10, 11, 12, 13]. The goal of CF is to impute missing preference elements in a sparse matrix consisting of users (rows) and items (columns), by solving a matrix completion task. The method predicts the unknown preferences for the users by elaborating correlations between known preferences of a group of users. Neural collaborative filtering (NCF) [14] is an advanced form of collaborative filtering that models the user-item interactions with a multi-layer feedforward neural network. CF models are commonly used to construct recommendation systems by companies like Netflix, Amazon, Linkedin, and Pandora [15], which provide popular benchmark datasets. Predicting drug activity for multiple targets can be formulated as a matrix completion problem analogous to a recommendation system, based on the credible assumptions that (1) there are a few dominant factors deciding which compound has which activities against a particular target (i.e., a low-rank approximation can be used) and (2) compounds with similar chemical structures will exhibit similar activities against similar protein targets. Compound activity prediction can be translated to a recommendation system problem by mapping compounds to users and targets to items. These methods work best when there are features (e.g., protein structures or sequences) that relate targets to one another. This presents a difficulty when using databases like DTC, ChEMBL, and ExcapeDB where target structures are usually not available. This also imposes limits on the types of properties used in CF approaches. Properties like synthesizability or solubility, which don’t relate to a particular protein target, do not fit well with the collaborative filtering paradigm.

### pQSAR 2.0

pQSAR 2.0 [16] uses properties, such as a IC_50_s on a panel of targets as features to build a model for compound induced activity inhibition for a target of interest. Martin et al. hypothesize that using these features are more informative than structurally derived features such as Morgan fingerprints [17]. For example, the muscarinic acetylcholine receptor (CHRM) family of proteins show highly correlated response values. It could be easier for a model to learn to predict CHRM1 inhibition using CHRM2-5 inhibition as a feature. Let *P* = {*p*_1_… *p_N_*} be an assembled panel of *N* targets. Let *C* = {*c*_1_… *c_M_*} be the set of *M* compounds from all assays for all targets. Assays are combined using the method described in the Data section.

The *MxN* matrix formed by compounds and targets is called a profile. A profile can be sparsely populated and sometimes has less than 2% of the cells filled. Random forest regressors (RFRs) are then trained for each individual target based on the existing data, using a conventional set of features such as chemical descriptors or fingerprints. All available data are used to train these RFRs producing *production RFRs.* These production models have seen *all* available data.

Next, partial least squares (PLS) models are built for each assay, using the measured values in the profile for the rest of the assays as features. The relationship between targets is learned by the PLS model; no target features are necessary. When the model is used for inference, missing values in the sparse profile are filled in with predictions from the RF models; these together with the measured values are used by the PLS models for the final activity predictions. This approach has been shown to outperform multitask models and, in our experiments, neural collaborative filtering as well (results not shown).

pQSAR has the added benefit of enabling new targets to be added easily. Each new target only requires training a new PLS model, which is significantly more computationally efficient than training a neural network (NN) model. This approach also lends itself to combining data from different sources, such as pharmaceutical companies, without having to explicitly share the proprietary structures [18].

The choice of assays to include in a profile affects performance [19]. Martin et al. choose assays for inclusion in a profile by training two models using two thresholds to filter models based on squared Pearson correlation (PR^2^) [20]. The higher scoring of the two models is recorded. In this work we seek to determine pQSARs performance before training models. Making this decision before processing large datasets with thousands of assays and millions of compounds can then be used to anticipate where pQSAR models are needed to improve property prediction performance.

Though pQSAR is very promising and has been shown to be very effective in several situations, there are specific conditions where pQSAR accuracy does not improve over single task models. Our application of pQSAR to GPCR and safety targets pulled from Drug Target Commons (DTC) [3], ExcapeDB [4], and ChEMBL [2] showed only 37 of 149 targets improving when compared against single task random forest regressors.

In this paper we compare pQSAR performance using production RFRs and pQSAR performance on actual compounds. pQSAR makes the assumption that compounds in the test set may have been used in the training set of the production RFRs. Evaluating pQSAR on virtual compounds means compounds in the test set must not appear in the training set for the RFRs. This makes a notable difference in prediction error. We train RFRs using just the training set and also production RFRs that use both training and test sets as described in [16]. Two pQSAR models are then built, one using RFRs and the other using production RFRs. These two pQSAR models are referred to as virtual pQSAR and actual pQSAR respectively. We seek to understand the conditions, which contribute to less favorable performance. Our work will elucidate the mechanisms behind pQSAR and help in finding new assays for inclusion in profiles. Heuristics developed in this paper will make it easier to build successful pQSAR models.

## Data

We obtained assay data for a combination of different targets and ligands from multiple data sources including Drug Target Commons (DTC) [3] (downloaded May 10th, 2020), ExcapeDB [4] (Version 1), and ChEMBL [2] (Version 27). Selected GPCR and safety assays as well as some safety assays were combined to form our profile (GPCR/Safety). GPCR binding activity data was obtained from ChEMBL 27 by first querying the database for human targets of type ‘single protein’ classified as ‘family A G protein-coupled receptor’. We used the resulting list of target IDs to select activities of type IC_50_, EC_50_, or Ki with units specified as nanomolar (nM), micromolar (uM), or ug/ml, and converted all measurements to nanomolar units. Outliers were filtered out, and replicate measurements were averaged to produce a single value of each activity type per compound per target. It is hard to know which experimental values to trust. Even after confirming identical assay procedures, variance in measured values is inevitable. As a result, the average of multiple measurements is used [21]. Target/activity type combinations with fewer than 200 compounds were dropped from the profile. The resulting profile contained 149 targets total and over 330,000 unique compounds. Assays with the same target identifier are combined across databases. All quantitative fields using uM, nM, ug/ml were converted to nM. Duplicates were averaged within one database, but removed in the case of inter-database collisions.

We also performed pQSAR with a profile comprised of kinase IC_50_s. This profile is comprised of 159 kinase assays downloaded from ChEMBL [2], on July 6 2015. The dataset was curated, split and previously published by Martin et al. [16] and is used as a benchmark.

## Results

Throughout this paper we compare pQSAR performance on two kinds of compounds. *Virtual compounds* refer to compounds that do not appear in any assays collected in this paper. Virtual compounds have no associated experimental data and only exist as a structure described by a SMILES string [22]. These compounds may appear in a drug design loop and only exist in a virtual sense. *Actual compounds* refer to compounds that appear in at least one assay collected in this paper. These compounds have been assayed against a target. Frequently we will pretend that an actual compound is a virtual compound by ignoring experimental data for the sake of validation.

### Multitask scaffold split

It is important to have a realistic split between training and test sets that accurately reflects the application of evaluating compounds of varying distances as defined in a chemical space. A randomized split usually produces higher model accuracy scores because it is more likely for compounds in the test set to be similar to compounds in the training set. In practice it is also desirable to evaluate structurally different compounds that are distinct from compounds used in the training set [16]. We chose to use the scaffold split implemented by DeepChem[23].

DeepChem’s scaffold split divides compounds into scaffolds as defined by Bemis et al [24]. For the data used in this study, this results in hundreds or thousands of scaffolds, most of which contain only one compound. Next scaffolds are assigned greedily by allotting the largest scaffolds to the training set until the training set is full. Then scaffolds are assigned to validation in the same manner. The validation set is a subset of the training data used to evaluate model performance during the process of selecting the best individual models. Any remaining scaffolds are assigned to the test set.

This method has two short comings. First, this is not the most optimal assignment of scaffolds with respect to average distance between sets in chemical space measured using Tanimoto distance [25]. Scaffolds can be shuffled between the three sets to make the average chemical distance between ligands larger. Second, in the multitask case, all labelled compounds for one task could be assigned to one set leaving no samples for the other two.

To address these two short comings we implement a novel splitting scheme that uses a genetic algorithm to assign scaffolds. This splitter divides the dataset into training, validation, and test subsets such that every subset for every task is populated while simultaneously maximizing the pairwise Tanimoto distance of compounds between subsets. Figure 1 demonstrates the difference using our multitask scaffold split described in Methods section, compared to a traditional single task split. This dataset was created by taking a subset of 10 tasks from ChEMBL. Directly comparing models trained using either split is difficult. A more rigorous split does not necessarily lead to a more generalizable model. In fact, a more rigorous split usually leads to a lower test score, which can be attributed to several variables. Instead, we measure the rigor of a split using pairwise Tanimoto distances between training and test sets.

**Figure 1.**
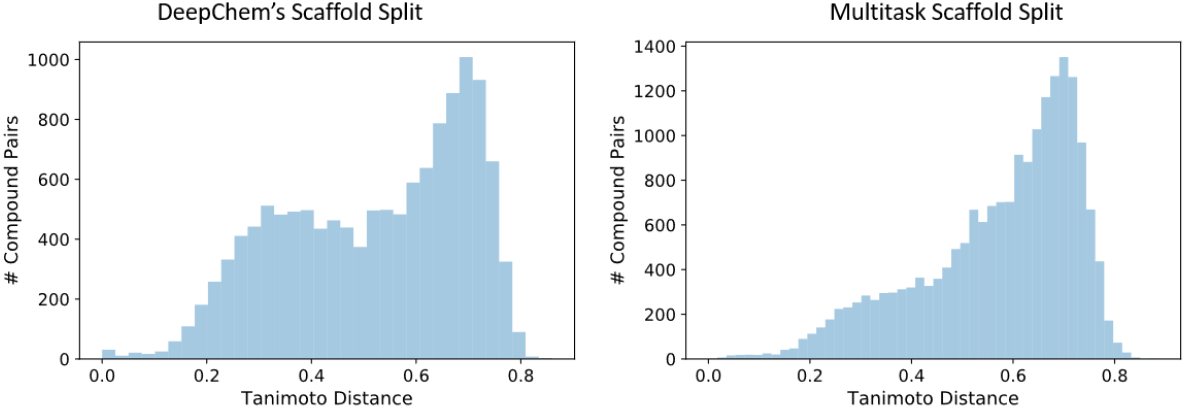
Left: The distribution of pair wise Tanimoto distances between training and test sets using DeepChem’s scaffold split. Right: The distribution of pair wise Tanimoto distances between training and test sets using our genetic algorithm, scaffold, split. The multitask scaffold split skews more to the right, meaning compounds are more different.

In Figure 1 we plot the distribution of pair wise distances between training and test sets. Using our genetic algorithm results in a distribution that skews more to the right, meaning compounds in training and test are further apart as measured by Tanimoto distance. The median pair wise distance is 0.55 using scaffold split and is 0.79 when using the genetic algorithm. In larger datasets, our 149 target GPCR/Safety profile for example, the change in distance distribution is less significant. However, while splitting our dataset, we found that using DeepChem’s scaffold split often resulted in some targets with no compounds in validation or test, typically not exceeding 3 targets. We use our genetic algorithm in these cases because the splits guarantee compounds in each set for each target and further justified using the genetic algorithm.

### Identifying ‘supported’ targets

We observe increased performance in pQSAR models when one or more of the following conditions are met. Targets that satisfy these conditions are referred to as *supported targets.* Targets that help satisfy conditions, but may not satisfy these conditions themselves are called *supporting targets.* Targets that do not satisfy any conditions and do not help satisfy conditions are referred to as *floating targets.* Our goal is to identify the set of conditions that allow pQSAR to succeed thus allowing users to evaluate their datasets before training anything.

We build a series of profiles to illustrate the effect of compound overlap between assays and profile fill percentage. First, we build a profile called GPCR/Safety Popular, that has a higher fill percentage than the original profile; see Table 1. Second, we look at the percentage of compounds in an assay that are used in at least one other assay, referred to as overlap percentage. This is slightly different from fill percentage, but is closely related. Adding an assay to the profile that shares compounds with another target can increase that targets overlap percentage, but lower the overall fill percentage. We create another profile called GPCR/Safety Support that contains all the targets in GPCR/Safety Popular plus additional assays that contain more overlapping compounds.

**Table 1.** Profile fill percentage

|  | Fill Percent |
| --- | --- |
| Kinase | 3.78% |
| GPCR/Safety profile | 1.00% |
| GPCR/Safety Popular | 1.68% |
| GPCR/Safety Support | 1.12% |

The most direct way to increase compound overlap is to increase the fill percentage of a profile, but this is not the only way to get improved models; see Table 1. The kinase profile has a fill percentage of 3.87%, significantly higher than our GPCR/Safety profile, which has fill percentage of only 1.00%. The GPCR/Safety Popular profile has a higher fill percentage than the GPCR/Safety profile, yet has fewer improved models; see Table 2.

**Table 2.** Task Info

|  | Total Tasks | Improved Tasks | Floating |
| --- | --- | --- | --- |
| Kinase | 159 | 151 | 9 |
| GPCR/Safety profile | 149 | 34 | 43 |
| GPCR/Safety Popular | 70 | 17 | 0 |
| GPCR/Safety Support | 106 | 26 | 0 |

GPCR/Safety Support contains all targets in the GPCR/Safety Popular profile and adds an additional 36 targets, 9 of which have pQSAR models that outperform single-task models. Of the 9 models that improved, 8 were newly added to the profile, and one was already present. Even though adding targets to the profile diluted the overall fill percentage, we were able to get more improved models.

Figure 2 contains bar plots for 4 profiles. Blue bars count the total number of targets that fall within the range of overlap percentage. Orange bars count the instances where pQSAR models outperform RFRs. We focus on the ratio of total targets to instances of improvement to determine when it might be beneficial to use pQSAR.

**Figure 2.**
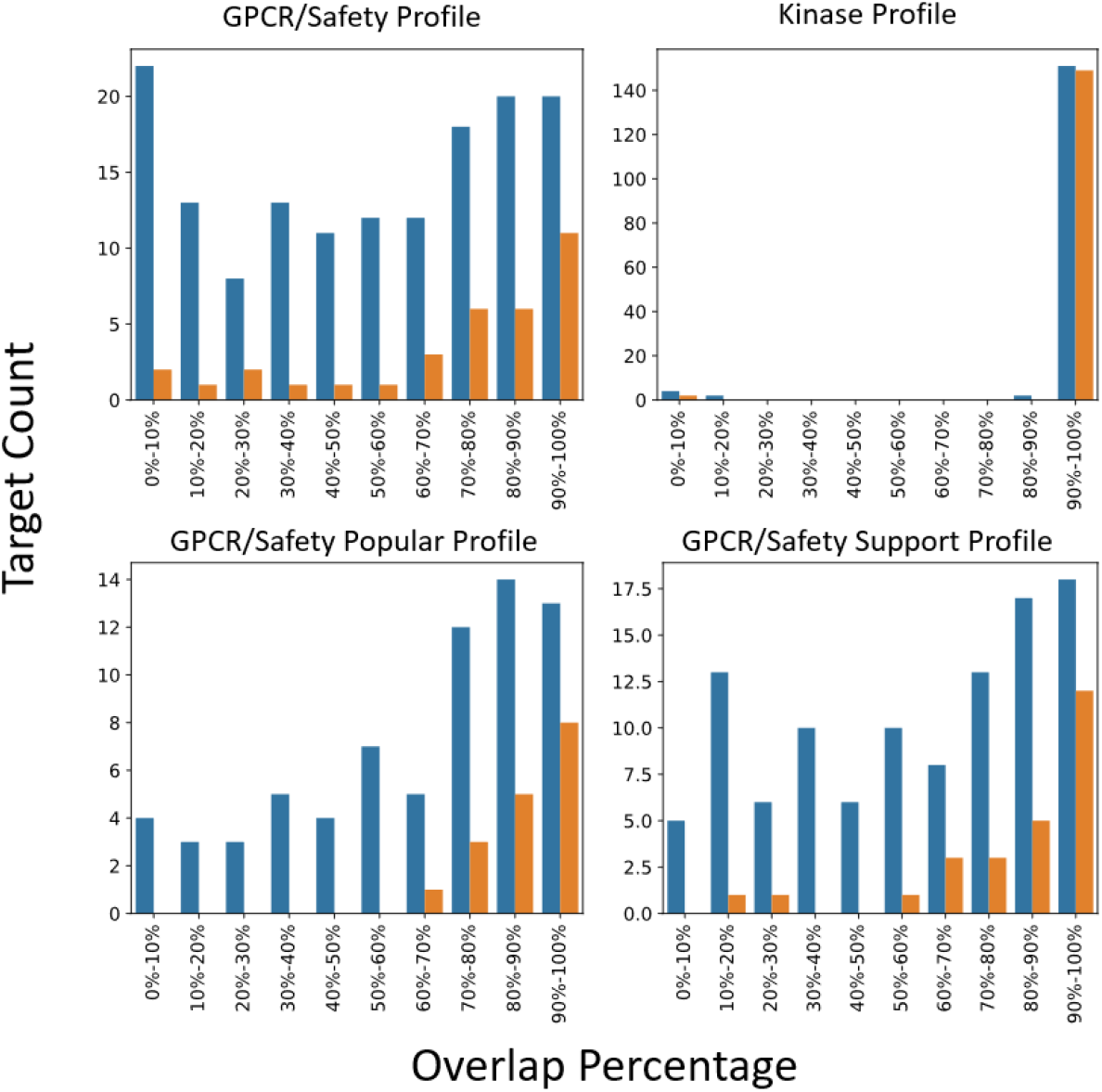
Blue bars indicate the number of targets that improved through the use of pQSAR. Orange bars indicate the total number of targets that fall in the range of overlap percentage. The higher the overlap percentage the higher the number of improved targets. Our observations lead us to believe that a minimum overlap percentage of .4 is require to motivate the application of pQSAR.

Targets that correlate with other targets in the profile will have more informative features to draw on during PLS training [19]. Linear models like PLS are not as expressive as higher dimensional models like SVMs, neural networks, or random forests. While this means that they are harder to overfit, it does also mean that features for PLS models must be informative and linearly related to the target. Many of the most successful targets in the GPCR/Safety profile belong to families or have other targets in the profile that strongly correlate (see Figure 3).

**Figure 3.**
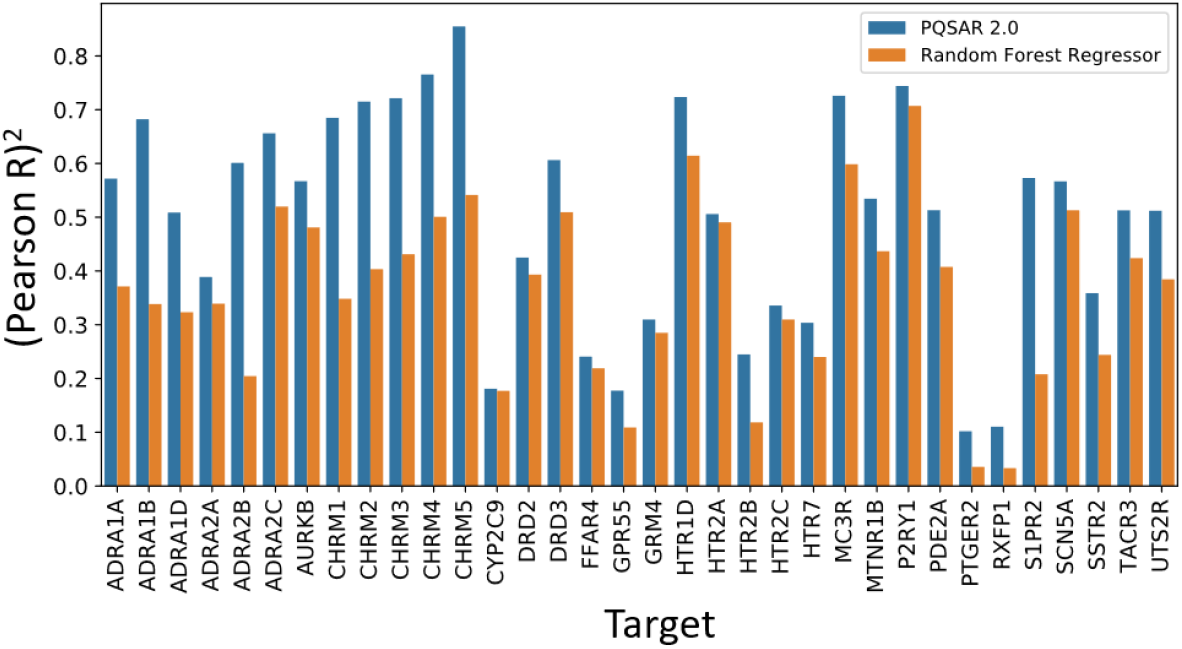
Comparison between RFR and pQSAR models for 34 tasks where pQSAR was superior. Most models that improved belong to families of closely related, highly overlapping targets.

### Virtual versus actual compounds

Figure 4a, reproduced from Martin et al. [16], plots the performance of random forest regressors (RFRs), pQSAR, and virtual pQSAR on the kinase profile. Targets are ranked by PR^2^ on the test set and plotted in decreasing order. The highest curve corresponds to pQSAR using production quality RFRs. This corresponds to the use case where one is predicting on actual compounds. In the other use case, virtual pQSAR performs poorly, but does improve on RFRs on some targets.

**Figure 4.**
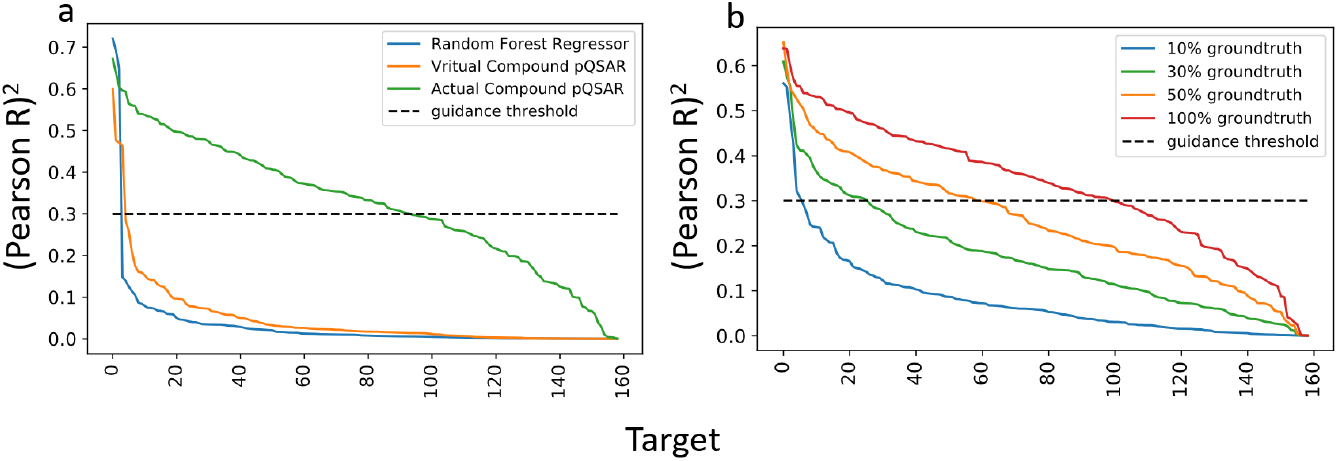
Performance curves for the kinase profile. a: Single task random forests perform very poorly on the test set. This translates to poor features for pQSAR which also performs poorly. Using pQSAR on actual compounds produces best results. b: pQSAR can produce good predictions on virtual compounds depending on the quality of single task classifiers.

Figure 4b shows results from restricting ground truth data. At 100% replacement, performance is nearly the same as the pQSAR model. As these replacements become less frequent, the model’s performance looks more and more like virtual pQSAR.

GPCR/Safety results shows a similar pattern in Figure 5a, where virtual pQSAR does not perform as well as pQSAR, though the performance gap between the two pQSAR models is smaller. While it looks like RFRs are superior to pQSAR, in fact there are 34 targets where pQSAR models have a higher PR^2^; see Figure 5b.

**Figure 5.**
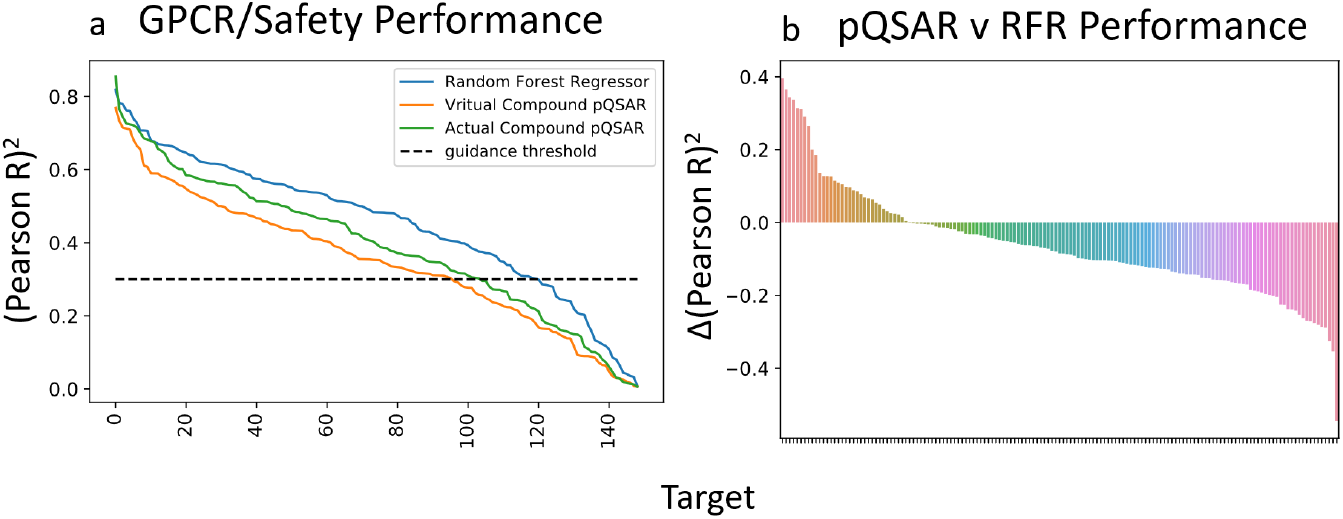
Results for GPCR/Safety profile. Left: performance of the pQSAR models on virtual compounds are lower than pQSAR models trained on production random forests. This time random forest models frequently out perform pQSAR. Right: pQSAR out performs random forests in some cases. This plot shows the difference in performance between the single task random forest and the production pQSAR.

### Hope for virtual compounds

Virtual compounds have not been seen during the training set and can be structurally distinct from compounds in the training set, thus presenting an especially challenging prediction problem for any data driven statistical model. Nevertheless, any potential information a prediction model can provide on virtual compounds is highly desirable since it would open a nearly unlimited number of new compounds to consider for their drug-like potential. For pQSAR to be applied to virtual compounds, the single task model predictions would need to meet a minimum prediction accuracy threshold. To investigate the minimum RFR accuracy requirements, predictions from actual compounds made by the production RFRs are modified to introduce varying amounts of noise to simulate the degradation in the RFR prediction accuracy and its impact on overall pQSAR performance. Details on the construction and training of these models can be found in the Methods section.

In Figure 6, we plot the results for several iterations of pQSAR on the kinase profile. In Figure 6a, we simulate a variety of RFR performances. The RF curve plots the performance of RFRs, evaluated on the test set, using only the training set. The production RF curve plots the performance of RFRs, evaluated on the test set, trained using both training and test sets. Curves between those two are made by adding Gaussian noise with zero mean and a given standard deviation to predictions made by production RFRs *if* the prediction is made for a compound in the test set. This allows us to create a series of models that will produce a prediction that is within one standard deviation 68% of the time. These simulated models are worse than a real model with the same performance on a given test set, because we know that the simulated models do not generalize well, Figure 4, where a real model would. Using these simulated models can give us a lower bound on how a pQSAR model would perform with models of varying accuracy.

**Figure 6.**
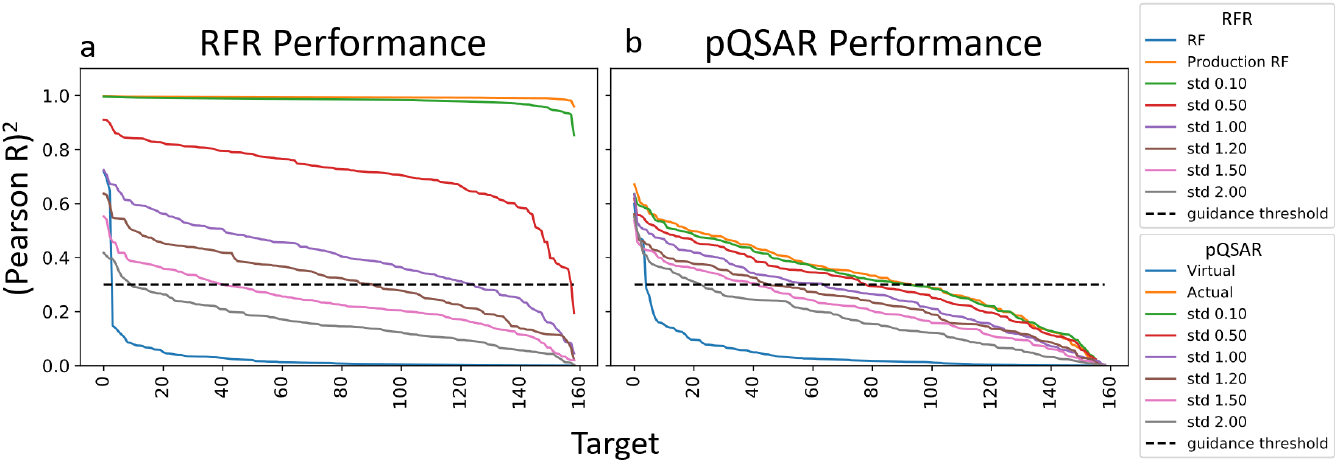
Increasing amounts of noise are added to ground truth values, left Profile, to simulate varying levels of performance in single task models. We can see a corresponding decrease in model performance, right pQSAR.

In Figure 6b pQSAR models are built using the simulated models as well as the two real RFR models. The Virtual pQSAR model is built using RFRs and the Actual pQSAR model is built using production RFRs. As noise increases, the performance curves drop, until the lowest curve which represents RFRs trained only on the training set. We observe that using RFRs with performance between 0.6 and 0.8 produces predictions nearly as good as the production RFRs. It is not until RFR performance drops below 0.4 that the pQSAR curves start to exceed performance. The simulation predicts that even RFR models with a standard deviation of up to 2 logs show pQSAR improvement in some targets, which indicates that pQSAR can be applied to virtual compounds in some cases. These predictions are verified in Figure 5 where the RFRs generalize well causing the gap between Virtual pQSAR and Actual pQSAR models to be smaller.

### pQSAR compared to single-task neural network models

We test the performance of 11 neural network models against the developed pQSAR models in order to compare prediction performance against other models representative of an extensive model training pipeline. Details of the construction of these models can be found in the methods section. The results are listed in Table 3. pQSAR improved the performance of modeling the CHRM assays. The PLS models for these assays had high coefficients (greater than 0.3) for at least two profile assays. Highly weighted assays were all from the CHRM family with the exception of adrenergic receptor beta (ADRB1), which was favored by the CHRM3 model with a coefficient of 0.241.

**Table 3.** pQSAR compared to single task NN models and RFRs. Scores are in PR^2^

| Task | Train # NN | Train # pQSAR | Train # RFR | NN | pQSAR | RFR |
| --- | --- | --- | --- | --- | --- | --- |
| AURKA | 4358 | 3302 | 2599 | **0.647** | 0.601 | 0.605 |
| AURKB | 3166 | 2607 | 1857 | **0.595** | 0.567 | 0.481 |
| CHRM1 | 1899 | 2727 | 2055 | 0.295 | **0.685** | 0.348 |
| CHRM2 | 1493 | 2121 | 1383 | 0.425 | **0.715** | 0.404 |
| CHRM3 | 1754 | 2444 | 1843 | 0.427 | **0.721** | 0.431 |
| CYP2C9 | 11321 | 10658 | 8233 | **0.209** | 0.181 | 0.177 |
| CYP3A4 | 10445 | 8571 | 5351 | **0.138** | 0.115 | 0.129 |
| CYP2D6 | 17796 | 15594 | 12189 | **0.389** | 0.159 | 0.317 |
| HRH1 | 1077 | 1925 | 1475 | **0.519** | 0.385 | 0.484 |
| KCNH2 | 11917 | 8193 | 5496 | **0.485** | 0.144 | 0.405 |
| PIK3CG | 2611 | 1466 | 1194 | 0.557 | 0.323 | **0.561** |

The PLS models for threonine-protein kinase aurora-A (AURKA) and threonine-protein kinase aurora-B (AURKB) used each other heavily, 0.624 and 0.834 respectively. These models did not weight other assays highly, with the majority of weights having a magnitude < 0.1. Coefficients for PLS models built for the cytochrome (CYP) family, histamine h1 receptor (HRH1), potassium voltage-gated channel subfamily H member 2 (KCNH2), and phosphoinositide-3-kinase catalytic gamma polypeptide (PIK3CG) also followed the pattern of having low weights. The highest observed weights for those assays was 0.220 in HRH1 looking at HRH2.

## Discussion

The kinase profile differs from the GPCR/Safety profile in notable ways that affects pQSAR performance. First, protein kinase inhibitors, by in large, work by displacing an ATP cofactor. This makes inhibitory affects of compounds highly correlated across different kinases. Second, assays created to screen for kinase inhibitors will frequently test the same compound across a panel of kinase targets. This can be observed in Figure 2 where almost all targets in the kinase profile have coverage between 90% to 100%. Together, the high correlation of activity and compound overlap make this the ideal target for pQSAR.

On the other hand, assays in the GPCR/Safety profile have nearly disjoint sets of compounds where most assays have less than 50% compound overlap. This profile contains a wider variety of targets collected without thought of correlation between targets, with the exception of some small families of highly correlated proteins. These targets all work with different mechanisms and compounds included across several targets are uncommon. This profile is the most ambitious case, trying to learn from nearly disjoint datasets, and stretches the limitations of pQSAR.

The PLS models rely on on high quality features generated by RFRs. In the absence of high quality RFRs, performance is determined by the presence of ground truth data, only available when predicting on actual compounds. This property can also be observed in the difference between pQSAR and virtual pQSAR in both kinase and GPCR/Safety profiles. RFR performance is high on GPCR/Safety targets, thus lowering the gap between the two pQSAR models.

GPCR/Safety Popular is built from a subset of targets that overlap frequently with other targets. These popular targets account for half of improved models in the GPCR/Safety panel, but the other half is left out of the profile. GPCR/Safety Support is built to include more targets that share compounds with targets in profile. These additional targets dilute the profile by adding compounds to the profile but also add targets with high compound overlap, which has a better chance of producing an improved pQSAR model.

From these profiles, we believe that a threshold of 40% makes the application of pQSAR worthwhile. This threshold is just a heuristic, as some models with lower overlap percentages also improve through pQSAR. This observation likely explains the increase in improved models when using the GPCR/Safety Support profile. The targets we added had high overlap percentages, thus adding more targets that had higher chance of success.

It is common for profiles generated in industry to have compound overlap [18], so limiting pQSAR to actual compounds from industrial compound libraries is not a set back. Additional steps should be taken when building profiles using disjoint assays and targets, whether that be collecting more assays for the profile, searching for more targets, or building better single-task models. We are in the process of building larger profiles, out of public data, to see if these observations continue to hold true.

In many cases, the majority of targets are not of interest, and one might care about just a handful of targets. The conditions defined in this paper can be combined with target sequence clustering [9] and pQSAR All-Assay-Max2 [19] to build profiles for maximizing performance of a subset of desirable targets.

Finally, using pQSAR for virtual drug screening is a very desirable application, and, as data scientists, we hope to be able to build models capable of replacing experimental measurements. These experiments were carried out using random forest regressors trained using Morgan fingerprints. Lenselink et al. trained a panel of machine learning models on targets from the ChEMBL database. Datasets were thresholded to transform a regression task into a classification task. To validate, they used a temporal split, which closely matches real world use cases. NN out performed support vector machines and random forests when validated on a temporal split [26]. There is some hope that more generalizable single-task models will lead to better virtual pQSAR models.

## Conclusions

Profile-QSAR provides a way to combine separate datasets into something that is more than the sum of its parts. Under the right conditions, correlated targets tested with overlapping compounds or access to accurate single task models, it is possible to use properties as independent variables to build a model on the target of interest. In our experiments with a profile built from the GPCR and safety targets, we saw an increase in performance in about 24% of targets.

Success of pQSAR boils down to compound overlap between assays and the amount of ground truth available for supporting targets. Assays that share a high percentage of compounds, greater than 40%, are more likely to have pQSAR models that out perform RFRs. This reliance on experimentally collected data is perhaps the weakest point in pQSAR and can be remedied with the improvement of the single-task models used to fill in the profile, in this case RFRs. Experiments we performed suggest that as the performance of RFRs or other models increase beyond a PR^2^ of above 0.6 to 0.8, it will be possible to use pQSAR models with virtual compounds. In addition, these heuristics could be used to prioritize future data collection efforts, where measurements on specific compounds or targets are anticipated to maximally improve predictive performance.

In future work, we will apply what is presented here to building profiles focused on specific targets of interest. By starting with a target, we can select assays from supporting targets that are correlated and share the same compounds. We believe that constructing profiles in this manner can be used to select the most predictive task specific model.

## Methods

### pQSAR 2.0

The *MxN* profile formed by M compounds and N targets is infilled using random forest regressors (RFRs) trained using Morgan fingerprints as input features. All the available experimental data is used to train these regressors.

Once features have been generated for each ligand, *N* partial least squares (PLS) models are fit to the target of interest, this time using only the training set. Each PLS model uses *P* – 1 features leaving out the *ith* column *p_i_* corresponding to the target of the PLS model. The results are compared to RFRs trained using only the training set, not the production RFRs that were used to generate features. This process is repeated for each target in the profile, building *P* PLS models.

Our implementation of pQSAR uses the ATOM Modeling PipeLine (AMPL) [27] to train random forest regressors (RFRs) and the Maestro [28] Workflow manager to manage each step of the process (e.g., data curation, RFR training, PLS training, tests, and visualizations). All processing was performed on a cluster containing 8-core Intel Xeon E5-2670 processors. RFRs were not limited in the number of estimators or depth to mimic decisions in [16]. A single RFR could take as long as 2 hours to train while it typically took 15 minutes to train all PLS models for a profile.

### Multitask scaffold split

Our multitask scaffold split uses a genetic algorithm to assign scaffolds to sets outlined in Figure 7. At each iteration we calculate the average distance between each pair of sets, (e.g. the average distance between training and test sets). Doing this on a per compound basis is intractable as is calculating pairwise distances between each scaffold. We group scaffolds into *S* super scaffolds to cut down on computation. Super scaffolds are created using the same method as training, validation, and test sets, now extended to S sets. This allows the genetic algorithm to reconstruct the same split as DeepChem if that split is optimal. The median distance between compounds in each pair of super scaffolds is saved in a distance matrix. The distance matrix is cached and used to calculate the distance between two super scaffolds.

**Figure 7.**
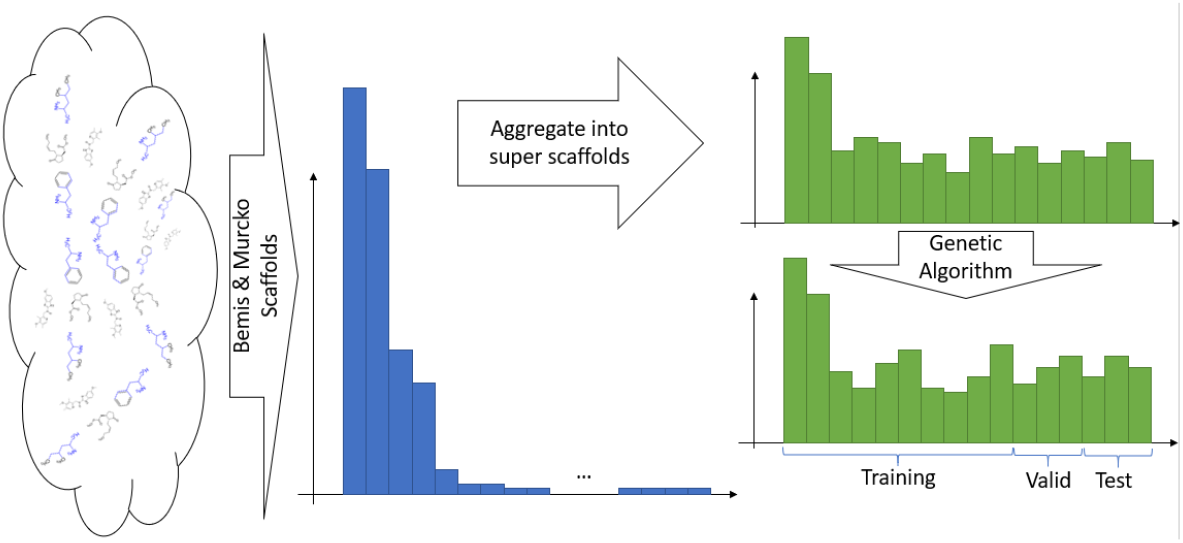
Outline of the Multitask Scaffold split algorithm. We aggregate scaffolds into super scaffolds and assign them to one of three sets using a genetic algorithm. The fitness score is a weighted sum of the chemical diversity between sets and how closely the sets matches the requested fractions.

A genetic algorithm then assigns each super scaffold to training, validation or test set. A chromosome is an array of S where each element in the array can hold one of three values train, validation, test. At each generation a chromosome’s health *H* is defined as:

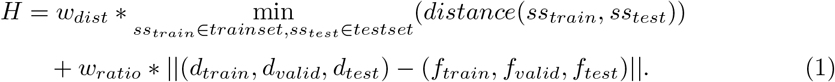

Where *ss_train_* is a super scaffold from the training set, *d_train_* is the desired training, validation, test fraction respectively, and *f_train_* is the actual training, validation, test fraction. The weights *w_dist_* and *w_ratio_* are tunable weights to balance the two terms. In our experiments we set both to 1. Any assignment that results in an empty set for a task is assigned a 0 health score.

The computational cost of our genetic algorithm split is significant. Spitting the 149 target GPCR/Safety takes hours longer to run, on an 8-core Intel Xeon E5-2670, relative to the single task splitter, which takes less than an hour on the same dataset. Fortunately, this split is done only once and can be cached for further experimentation.

### Profile Construction

GPCR/Safety Popular, contains a higher fill percentage than the original profile; see Table 1. Figure 8 shows how many compounds two targets have in common and the Spearman correlation between the two. Each column and row represents a target and are ordered on the number of non zero values in each column. We set the threshold at 70, which appears to be a natural drop off.

**Figure 8.**
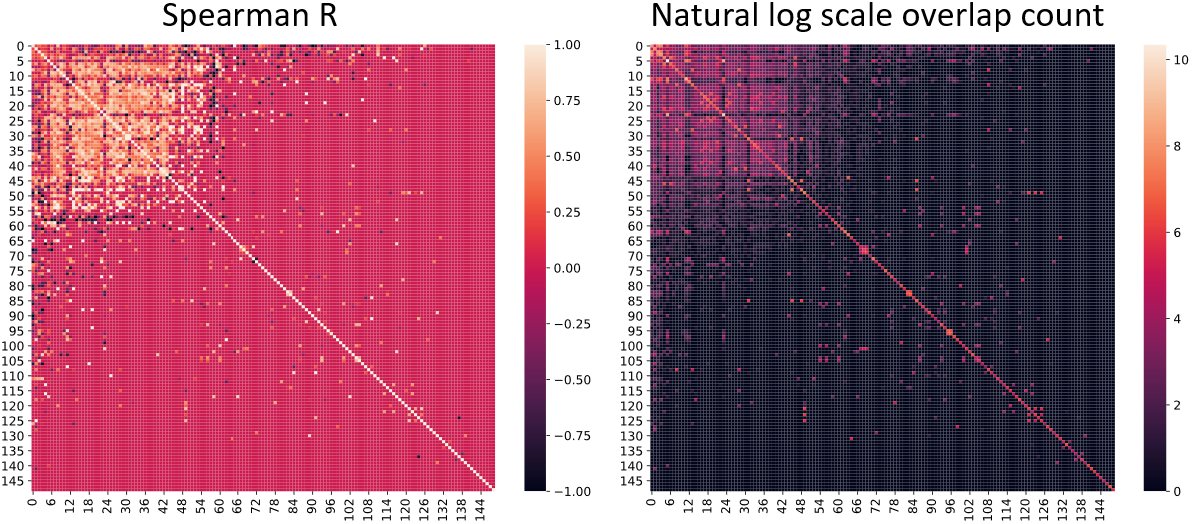
Heatmaps of cross Spearman correlation (left) and cross target ligand overlap on a natural log scale (right). The targets are ordered on the number of other targets that contain overlapping ligands, in other words, non zero entries in the matrix. There is a strong distinction between targets that frequently overlap with other targets and targets that do not share ligands.

GPCR/Safety Support contains all the targets in GPCR/Safety Popular plus additional assays that contain more than 50 overlapping compounds and correlate with Spearman correlation greater than 0.3. We use Spearman rank correlation instead of Pearson or correlation coefficient r^2^ because we wish to be able to extend pQSAR to include properties with different units, though all profiles in our experiments have the same units, nM. Figure 9 visualizes all selected targets and the amount of overlap between the targets on a log scale. Large clusters along the diagonal correspond to families of targets that share many compounds.

**Figure 9.**
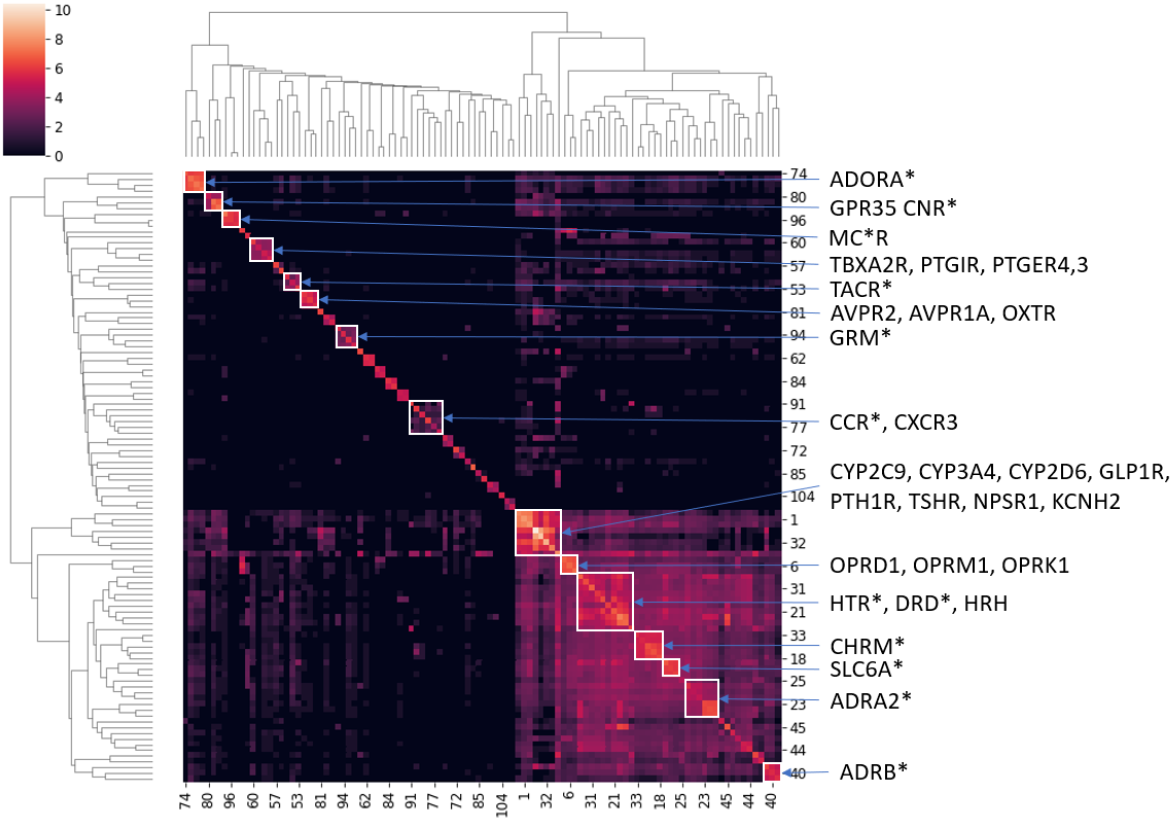
Heatmap of compound overlap between all tasks in GPCR/Safety Support. This visualization shows the selected tasks all overlap and support another task in some way. Clusters of targets are highlighted.

#### Examples

In Table 4, target A is a supported target. Supporting targets, like targets B, C, and D, overlap with supported targets, but may not have many overlapping compounds otherwise. Finally, floating targets have little overlap with other targets. We expect supported targets to have a boost in performance after applying pQSAR while supporting targets and floating targets are less likely to see any improvement. In this paper supported targets refer to the top 70 most overlapped and highly correlated targets. Supporting targets share at least 50 compounds with a supported target.

**Table 4.** Supported vs Supporting vs Floating.

|  | A | B | C | D | E |
| --- | --- | --- | --- | --- | --- |
| A | 300 | 50 | 100 | 70 | 2 |
| B | 50 | 100 | 2 | 10 | 0 |
| C | 100 | 2 | 200 | 4 | 0 |
| D | 70 | 10 | 4 | 150 | 10 |
| E | 2 | 0 | 0 | 10 | 300 |

### Virtual versus actual compounds

We build two types of pQSAR models using production RFRs and RFRs trained only on the training set referred to as actual pQSAR models and virtual pQSAR models respectively. Comparing the two allows us to measure the performance between evaluating compounds that appear in other assays and exploring totally new compounds not tested in any other assays.

We also experiment with restricting the amount of ground truth information in the profile to simulate increasing sparsity of data and how well pQSAR performs on compounds tested in fewer assays. First we construct a virtual pQSAR model, then whenever possible, we replace predicted values with ground truth data. We slowly scale back frequency of replacement, first to a frequency of 0.5 and then lower.

### Single-task neural network models

We also compare our models against pre-trained, single-task models for 11 safety assays that consider additional non-RF modeling approaches. These models followed a different pipeline and experimented using graph [29] and Mordred features [30]. Raw datasets for the 11 protein disease and safety related targets were downloaded from public data sources: Drug Target Commons (DTC) [3], ChEMBL [2], and ExcapeDB [4]. PIC_50_ for replicates were averaged or drop compounds that have large variances between their replicates. Then the data was split using DeepChem’s scaffold split into training, validation, and test sets. Data from the three sources were also combine to form a union training set and a union test set to train union machine learning models. Extensive comparisons between source specific models and union models showed that the machine learning models trained using the union training set performed best on both source-specific test sets and union test sets for most of the 11 targets. These experiments were made possible by using the AMPL open-source software package [27] which is built on DeepChem. Using AMPL it was possible to automate the training and comparisons of models using different hyperparameter settings including molecular descriptors used and machine learning methods e.g., random forests, neural networks, and XGBoost. Previous work has shown that this procedure leads to high quality models [31].

Since these models were developed using a different pipeline, special consideration needed to be taken when comparing prediction results. A special test set was built using the difference between the test set from pQSAR experiments and the union training set used in training the single task neural network models. After removing compounds seen in the union train set, each of the targets still contained at least 100 samples except PI3-kinase class I (PIK3CG) which only contained 64 samples.

## Acknowledgements

We thank Accelerating Therapeutics for Opportunities in Medicine (ATOM) for their support. We also thank Eric Martin for his advice. Thank you to Warren He who helped edit this manuscript. This work was performed under the auspices of the U.S. Department of Energy by Lawrence Livermore National Laboratory under Contract DE-AC52-07NA27344

This document was prepared as an account of work sponsored by an agency of the United States government. Neither the United States government nor Lawrence Livermore National Security, LLC, nor any of their employees makes any warranty, expressed or implied, or assumes any legal liability or responsibility for the accuracy, completeness, or usefulness of any information, apparatus, product, or process disclosed, or represents that its use would not infringe privately owned rights. Reference herein to any specific commercial product, process, or service by trade name, trademark, manufacturer, or otherwise does not necessarily constitute or imply its endorsement, recommendation, or favoring by the United States government or Lawrence Livermore National Security, LLC. The views and opinions of authors expressed herein do not necessarily state or reflect those of the United States government or Lawrence Livermore National Security, LLC, and shall not be used for advertising or product endorsement purposes. LLNL-JRNL-821120-DRAFT

## Abbreviations

GPCR: G protein-coupled receptor
pQSAR: profile - quantitative structure-activity relationship

## Availability of data and materials

We will release the data curation code on GitHub: https://github.com/ATOMconsortium/AMPL.

## Ethics approval and consent to participate

Not applicable.

## Consent for publication

All authors give their consent for this to be published..

## Authors’ contributions

SH: Helped writing the software and was the lead in writing the paper. SK: Helped writing software and edited the paper. KSM: Advised writing the software, edited the paper, and helped curate datasets. HR: Trained single-task neural networks and helped curate datasets. JEA: Advised writing the software, curated datasets, and edited the paper. DS: Helped edit the paper and curated datasets.

